# Synergistic Integration of Deep Neural Networks and Finite Element Method with Applications for Biomechanical Analysis of Human Aorta

**DOI:** 10.1101/2023.04.03.535423

**Authors:** Liang Liang, Minliang Liu, John Elefteriades, Wei Sun

## Abstract

Motivation: Patient-specific finite element analysis (FEA) has the potential to aid in the prognosis of cardiovascular diseases by providing accurate stress and deformation analysis in various scenarios. It is known that patient-specific FEA is time-consuming and unsuitable for time-sensitive clinical applications. To mitigate this challenge, machine learning (ML) techniques, including deep neural networks (DNNs), have been developed to construct fast FEA surrogates. However, due to the data-driven nature of these ML models, they may not generalize well on new data, leading to unacceptable errors.

**Methods:** We propose a synergistic integration of DNNs and finite element method (FEM) to overcome each other’s limitations. We demonstrated this novel integrative strategy in forward and inverse problems. For the forward problem, we developed DNNs using state-of-the-art architectures, and DNN outputs were then refined by FEM to ensure accuracy. For the inverse problem of heterogeneous material parameter identification, our method employs a DNN as regularization for the inverse analysis process to avoid erroneous material parameter distribution.

**Results:** We tested our methods on biomechanical analysis of the human aorta. For the forward problem, the DNN-only models yielded acceptable stress errors in majority of test cases; yet, for some test cases that could be out of the training distribution (OOD), the peak stress errors were larger than 50%. The DNN-FEM integration eliminated the large errors for these OOD cases. Moreover, the DNN-FEM integration was magnitudes faster than the FEM-only approach. For the inverse problem, the FEM-only inverse method led to errors larger than 50%, and our DNN-FEM integration significantly improved performance on the inverse problem with errors less than 1%.

## 1. Introduction

Machine learning (ML), especially deep (machine) learning using deep neural networks (DNNs) [1], has the potential to transform the biomedical and healthcare fields by enabling more personalized and accurate diagnoses and prognoses, treatment recommendations, and disease prevention strategies [2]. For example, in the cardiovascular domain, ML techniques have been developed for various applications, such as automated ECG signal analysis [3], automated geometry reconstruction from medical images [4], tissue modeling [5-8], heart valve analysis [9], disease modeling and diagnosis [10-12].

Almost parallel to the evolution of ML techniques, finite element method (FEM) has been established as a standard tool for mechanical analyses in many engineering problems, including structural analysis of human tissues and organs. For example, by using FEM to solve nonlinear large deformation, the stress and deformation of human aorta can be obtained under different physiological conditions (e.g., normal blood pressure vs hypertension), and the analysis results could offer biomechanical insights into various disease conditions, such as rupture/dissection risk of aortic aneurysm [13, 14]. Therefore, finite element analysis (FEA, analysis using FEM) on a patient-specific level have been widely explored for its utility in the diagnosis and prognosis of cardiovascular diseases. However, patient-specific FEA needs extensive human-user interactions and long computing time, which makes it unsuitable for time-sensitive applications such as rupture/dissection risk assessment of aortic aneurysm. To overcome this limitation, in our previous studies, we demonstrated the feasibility of using data-driven ML surrogate models for fast biomechanical analysis [15-18]. For instance, we developed a DNN model as fast surrogate of FEA for stress analysis of the aorta [16], which marks the first work that employed DNNs to establish an end-to-end nonlinear mapping between 3D geometry and 3D stress field.

Although ML has great potential and has made significant achievements, there are several limitations and challenges, such as bias, interpretability, robustness, and generalization, which must be considered when using ML for life-critical and time sensitive clinical applications. The fundamental limitation/challenge is the generalization capability of an ML model, which is the ability of an ML model to make the right decisions for new samples that are unseen during training. Based on the universal approximation theorems (UATs) [19], a DNN can approximate any continuous function with arbitrarily small error, if provided with enough capacity. UATs explain that a DNN can reach close-to-zero error on training data, but excellent performance of a DNN on test samples is not theoretically warranted. As will be shown in this study, a trained DNN can predict aorta deformation and stress with small/acceptable errors in majority of test cases; yet, for some test cases that could be out of the training distribution (OOD), the errors in the predicted peak stress were larger than 50%. For life-critical clinical applications, it is highly demanded that a DNN can guarantee or at least know its accuracy for any input sample. However, it remains challenging to estimate the accuracy of a DNN without making simplifications and assumptions [20].

In this work, we propose a synergistic integration of DNNs and FEM to overcome each other’s limitations and demonstrate this approach for the forward and inverse analysis problems of human aorta biomechanics. For the forward problem, we develop five types of DNNs for deformation prediction and combine each of the DNNs with FEM. We show that the integrated DNN-FEM is capable of novelty detection (i.e., whether the input is an outlier) and makes fast prediction with guaranteed accuracy, and it runs magnitudes faster than the FEM-only approach. For the inverse problem, we develop a DNN-FEM inverse method and show its advantage over a FEM-only inverse method for heterogeneous material parameter identification.

### 2. Methods

## 2.1 Data for the forward problem

The forward problem is to predict the deformed geometry of the aortic wall at the systolic pressure, for which a zero-pressure geometry and material parameters are given as the inputs, and the inlet and outlet boundary nodes of the aortic wall are fixed with zero-displacement. Here, a zero-pressure geometry refers to the geometry of an aorta at zero blood pressure level, and it is considered deformation/stress-free by ignoring residual deformation/stress. Once the deformed geometry is obtained, the stress on the wall can be calculated by using the deformation and the constitutive model of the tissue. To demonstrate our approach, we used a statistical shape model and FEM to generate numerical cases so that the ground truth (i.e., true deformation, true stress, and true material parameters) is available for DNN training and evaluation.

To model the hyperelastic behavior of aorta tissue, we used the Gasser-Ogden-Holzapfel (GOH) constitutive model [21]:

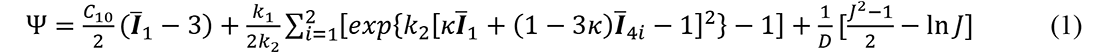

The GOH model has a set of material parameters {*C*_10_, *k*_1_, *K*_2_, *κ*, *θ*}. *Ī*_1_ and *Ī*_4i_ are strain invariants. *D* is a constant to enforce material incompressibility. Given the material model in Eq.(1) and the deformation gradient tensor ***F***, the Cauchy stress tensor ***σ*** can be calculated by

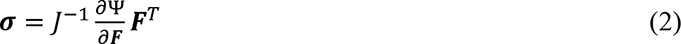

where *J* = det (***F***). The deformation gradient tensor ***F*** can be calculated by using the zero-pressure geometry and the deformed geometry.

We created a dataset of 342 zero-pressure geometries using a statistical shape model (SSM) that is built on 60 real aorta geometries of thoracic aortic aneurysm patients, and we created 125 sets of material parameters by sampling from the material parameters of aortic wall tissues. We refer the reader to our previous work for the detailed procedures of building a SSM and sampling material parameters [5, 15]. Each of the 342 zero-pressure geometries is paired with each of the 125 material parameter sets, and then it is inflated by using FEM with the systolic pressure that is fixed to 18KPa in this study, representing hypertension stage-1 [22]. An aorta geometry is represented by a hexahedron mesh of 10000 nodes and 4950 elements.

We conducted two types of numerical experiments for the forward problem. In the experiment-1, we use a representative set of material parameters for data generation, and then train and test DNNs to predict the deformed geometry at systole, given a zero-pressure geometry as input. In the experiment-1, the whole dataset is divided into a training set (171 samples) including 50% of the shapes, a validation set (34 samples) including 10% of the shapes, and a test set (137 samples) including 40% of the shapes. In the experiment-2, we use all the material sets and shapes, choose the best-performed DNN in the experiment-1, and then train and test the DNN to predict the deformed geometry at systole, given a zero-pressure geometry and material parameters as inputs. Given the deformation prediction from a DNN, the stress is computed by using Eq.(2). In the experiment-2, the whole dataset is divided into a training set (17100 samples) including 50% of the shapes and 80% of the material sets, a validation set (408 samples) including 10% of the shapes and 10% of the material sets, and a test set (1781 samples) including 40% of the shapes and 10% of the material sets. For each experiment, the training, validation, and test sets do not share any shapes nor materials.

## 2.2 DNNs for deformation prediction

We developed five types of DNNs to predict the deformed geometry of an aorta for the forward problem, which are MLP, W-Net, TransNet, U-Net, and MeshGraphNet. Here, MLP refers to multilayer perceptron, serving as the baseline model. W-Net is a brand-new architecture designed by us in this study. We designed TransNet by borrowing the Transformer architecture [23] that was originally proposed for nature language processing (NLP). U-Net [24] is widely used for image analysis, and we modified it for this study. MeshGraphNet [25] was proposed for mesh-based simulation, and we adapted it for this study. In the experiment-1, we performed grid-search to find optimal DNN configurations using the training and validation sets, and we refer the reader to the Appendix for more details. The best DNN in the experiment-1 is W-Net, and therefore only W-Net is chosen for the experiment-2.

### 2.2.1 MLP encoder-decoder for deformation prediction

The simplest DNN for this task is an MLP that has an encoder and a decoder. The encoder compresses the input, i.e., a zero-pressure geometry denoted by *X* = {*X*[*n*], *n* = 1,2, …} where *n* is node index, from a high dimensional space into a code vector *c* = {*c*[*i*], *i* = 1,2, …} in a much lower dimension. The decoder transforms the code into the final output, i.e., the predicted displacement field 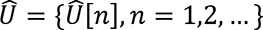. Then the deformed geometry 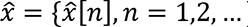 is obtained by 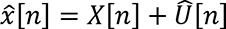. By using the Eq.(2) with X and 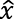, the stress at each element can be calculated.

### 2.2.2 W-Net for deformation prediction

We designed W-Net, a weight-constrained network whose internal weights are generated and constrained by sub-networks. As shown in Figure 1, for the experiment-1, the W-Net has an encoder-decoder structure, which is defined by the following equations:

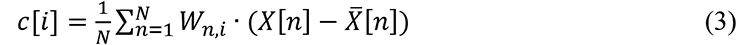

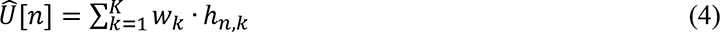

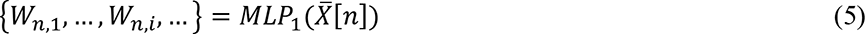

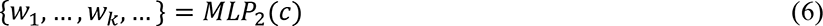

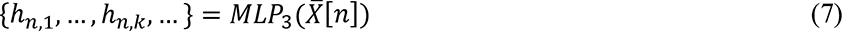

**Figure 1.**
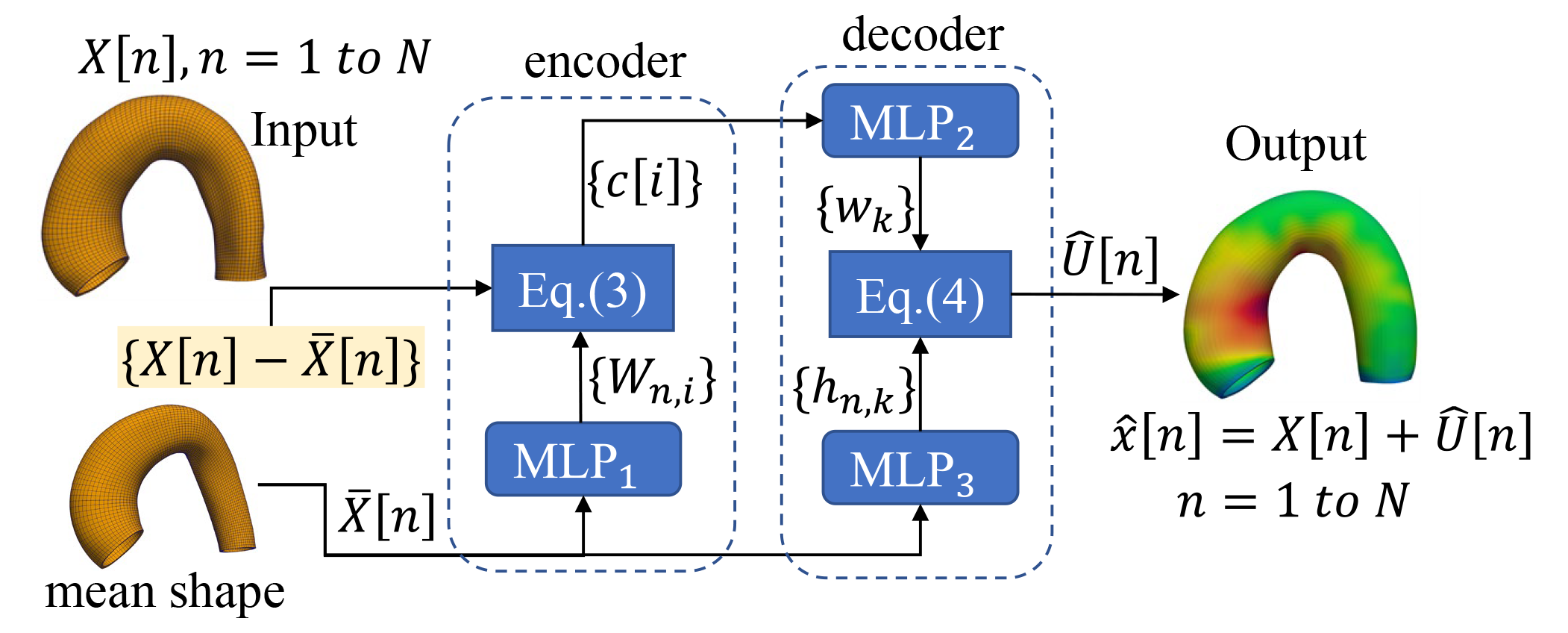
The structure of W-Net to predict the deformation.

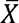 denotes the mean shape of the training samples, and 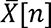 denotes the 3D coordinates (a vector) of the node-*n* on the mean shape. Similarly, *X*[*n*] denotes the 3D coordinates of the node-*n* on the mesh of an aorta at the zero-pressure state, and 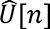 is the 3D displacement of the node-*n*. *N* is the total number of nodes on a mesh. *c* is the code vector, and *c*[*i*] is the *i*-th element of the code vector. *W*_*n*,*i*_ is a weight vector generated by an MLP with 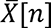 as input. *w_k_* is a weight scalar generated by an MLP with the code vector *c* as input. *h*_*n*_,_*k*_ is a vector generated by an MLP with 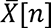 as input.

For the experiment-2, the inputs of W-Net include not only a zero-pressure geometry but also material parameters. To handle the material input, we first apply an MLP to encode the material parameters:

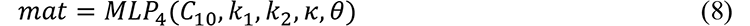

Then, we concatenate the original code vector and the new vector *mat* to obtain a new code vector by *c* ← [*c*, *mat*].

### 2.2.2 TransNet for deformation prediction

We designed TransNet for deformation prediction using the Transformer as the backbone. The Transformer is a neural network architecture originally introduced in the field of natural language processing (NLP) [23]. It consists of an encoder and a decoder, both of which use multi-head self-attention mechanism that allows the model to identify important context for each word. Since its introduction, the Transformer has become the backbone of many state-of-the-art NLP models, such as ChatGPT. In addition to its success in NLP, the Transformer architecture has also been applied to other domains, such as computer vision [26], and molecular and biological sciences [27].

The self-attention mechanism in the Transformer needs to calculate the attention matrix given by:

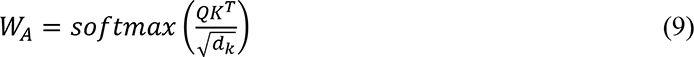

*Q* is called query, in which the number of rows equals to the number of tokens (e.g., a word in a sentence for NLP), and the number of columns equals to the number of features denoted by *d*_*k*_. *K* is called key, and it has the same size as Q. Each row of Q is a function of a token, and the same is true for *K*. A naïve implementation would be using every node of an aorta mesh as a token and calculating the self-attention matrix *W*_*A*_ among those nodes, which will result in a 10000× 10000 matrix *W*_*A*_. This naïve implementation with a few attention heads and layers requires huge amount of GPU memory that is not available even in a high-end GPU (e.g., Nvidia RTX A6000).

The structure of the TransNet is shown in Figure 2. The mesh of the aorta consists of circumferential curves, and each curve consists of 100 nodes. In total, an aorta mesh has 100 circumferential curves, circling around the centerline. We treat a circumferential curve as a token and obtain its embedding by using a linear projection as the circumferential curve encoder, for which the inputs are the coordinates of the 100 nodes on the curve-*m*, and the output is its embedding (a row vector) denoted by *h*_*m*_. We compute the centerline of the mean shape and apply an MLP to obtain the embeddings of the 100 centerline points, denoted by {*β*_1_, … *β*_*m*_, … *β*_100_} which contain the relative position information of the circumferential curves. To encode the position information into an embedding *h*_*m*_, we applied the position-embedding operation: *h*_*m*_ ← *h*_*m*_ + *β*_*m*_. The current embeddings {*h*_*m*_} (a.k.a. feature vectors) of the tokens will go through a sequence of multi-head self-attention layers, and a layer will output new embeddings of the tokens, which will be the inputs to the next layer. The last multi-head self-attention layer will output new embeddings of the 100 tokens, each of which corresponds to a circumferential curve. Given each of the final embeddings as the input, a linear transform is used as the circumferential curve decoder to predict the displacements of the 100 nodes on the curve from the zero-pressure state to the systolic phase. We refer the reader to the reference [23] for the details of the multi-head self-attention layer that is a basic component of Transformer.

**Figure 2.**
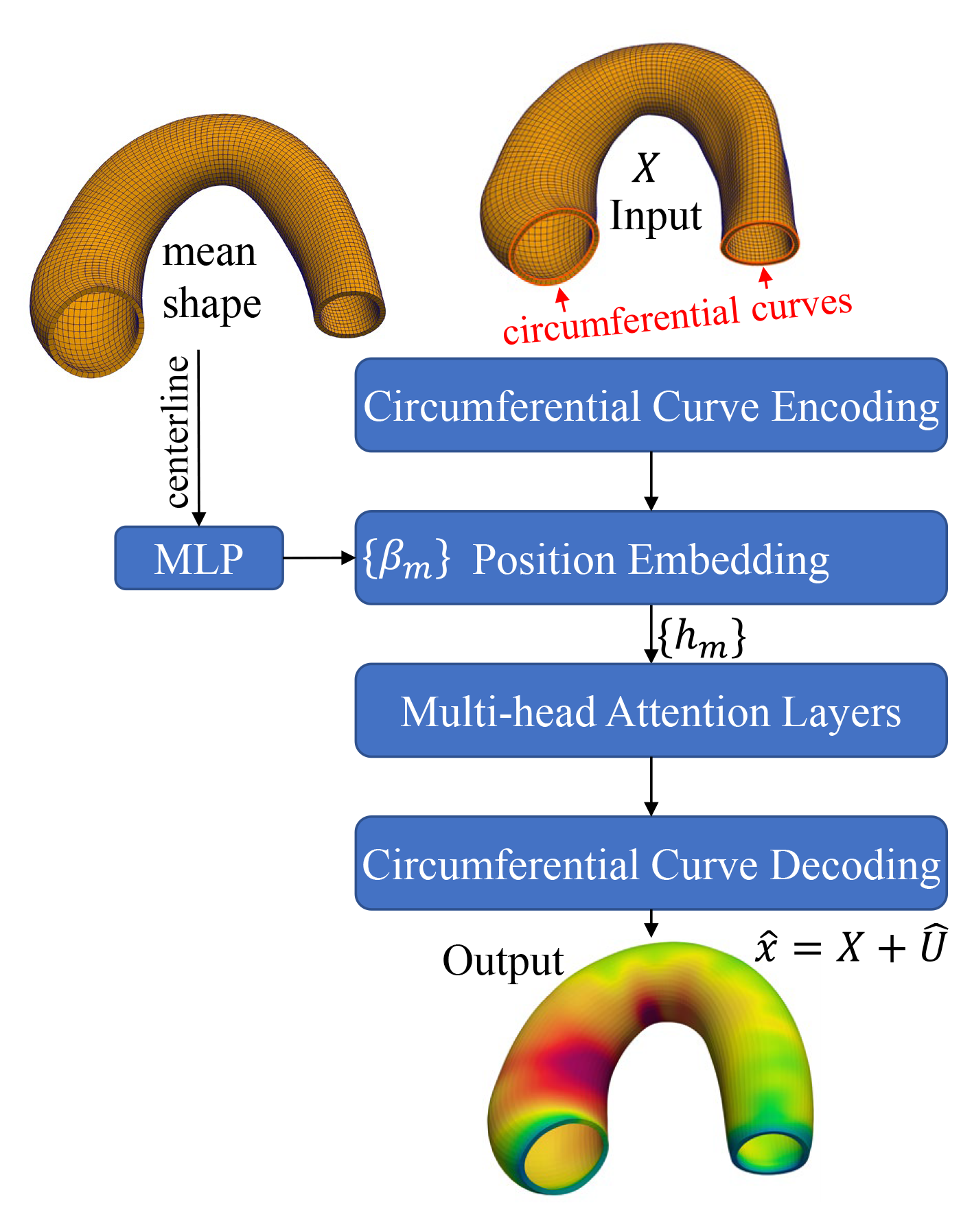
The structure of TransNet for deformation prediction.

### 2.2.3 U-Net for deformation prediction

We developed a U-Net with 1D convolution for deformation prediction, as shown in Figure 3. U-Net is a convolutional neural network architecture that was originally proposed for medical image segmentation [24]. It is named "U-Net" because of its U-shaped architecture with an encoder, a decoder, and skip connections, which integrates local and global information in a hierarchal fashion.

**Figure 3.**
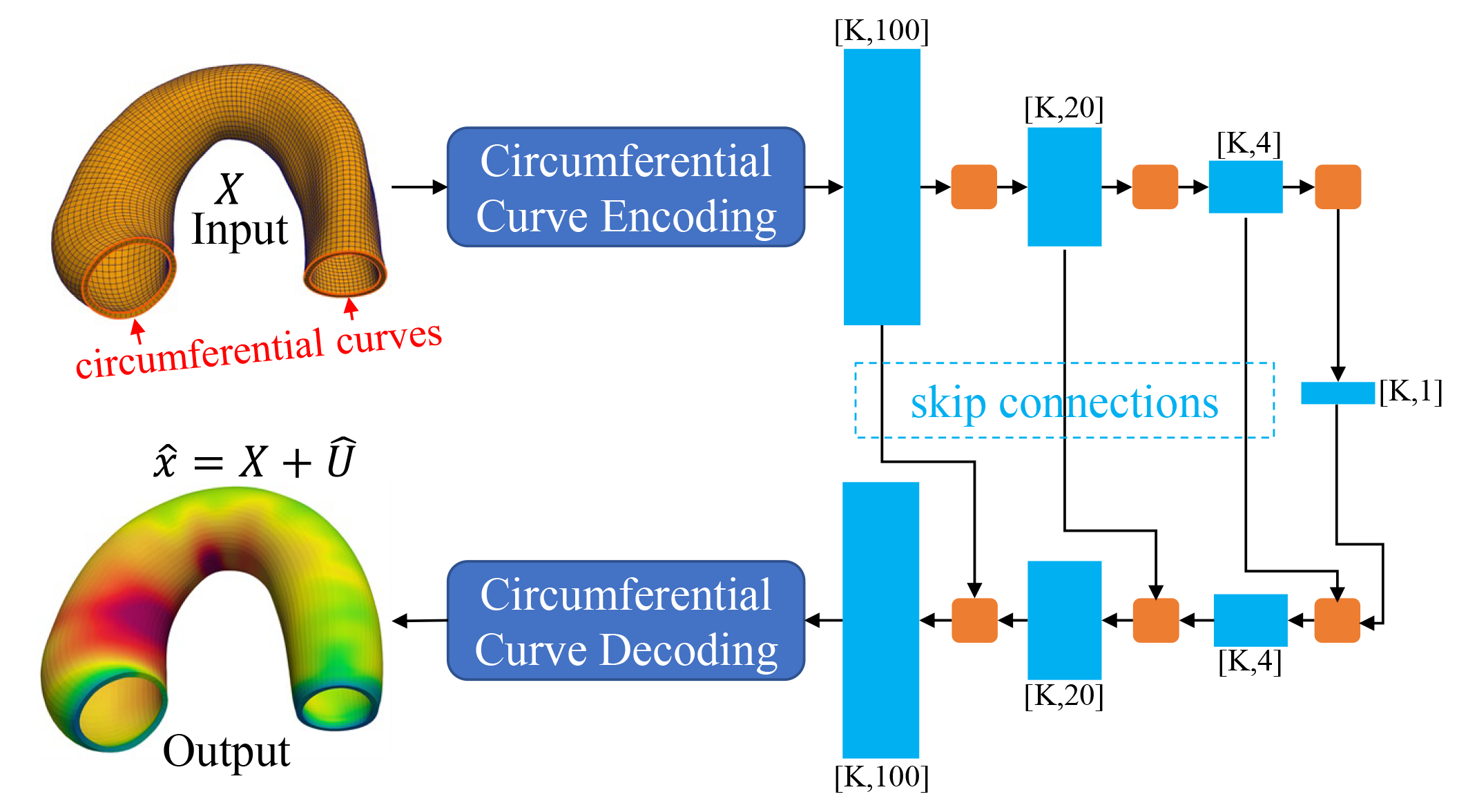
The structure of U-Net for deformation prediction. See the appendix for the value of K.

A circumferential curve of an aorta mesh is considered as a single pseudo point. An aorta mesh consists of 100 pseudo points aligned along the centerline; and each pseudo point has an embedding vector obtained by using an MLP as the circumferential curve encoder. Thus, the input to the U-Net encoder is a 1D sequence of 100 pseudo points, and the output from the U-Net decoder is a 1D sequence of 100 embedding vectors. Given each of the final embeddings as the input, an MLP is used as the circumferential curve decoder to predict the displacements of the 100 nodes on the curve from the zero-pressure state to the systolic phase.

### 2.2.4 MeshGraphNet for deformation prediction

MeshGraphNet [25] was designed for learning mesh-based simulation using a graph neural network. The network has an encoder to encode mesh information at each node, a processor to exchange information between the nodes, and a decoder to predict the displacement of each node. It can simulate deformable objects in a visually realistic manner.

In this study, we adapted this network to predict the displacement field of aorta from the zero-pressure state to the systolic phase. Compared to the other DNNs, in the MeshGraphNet, each node only has local neighbor information, and this information constraint is most likely the main reason that MeshGraphNet had relatively weak performance, as will be shown in the result section.

## 2.3 DNN-FEM integration for deformation prediction

We use the term DNN-FEM to refer to the integration of a DNN and FEM. As will be shown in the result section, even though the W-Net achieves the best performance, it may still make mistakes: the output errors are unacceptably large (>50% error in peak stress) for some test samples. For medical applications, it is expected that a model can guarantee or at least know the accuracy of its output.

In this study, we use FEM to refine the output from a DNN, so that the accuracy is guaranteed by FEM if the refinement process runs successfully, and the time cost of the refinement is minimal compared to a FEM-only approach that starts from the zero pressure state with numerous increments to gradually inflate the aorta mesh model to the systolic phase. To facilitate the integration of DNN and FEM, we implemented FEM using the basic functions in PyTorch, an open-source machine learning (ML) library. We refer the reader to our technical report about the implementation details and comparison with Abaqus [28], which shows that the discrepancy between our FEM implementation and Abaqus is negligible. Since both DNNs and FEM are implemented by using the same ML library, the integration can be made seamless.

The integration of a DNN and FEM is straightforward: the output displacement field 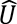 from a DNN will serve as the starting point (i.e., initial solution) of FEM to solve the forward problem. The issue is that the Newton– Raphson-based FE solver often crashes if 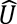 from a DNN is directly used as the starting point. To explain this issue, we provide a brief discussion here. Let Π represent the total potential energy as a function of the displacement field. From the perspective of optimization, the forward problem could be explained by using the principle of minimum total potential energy. By applying the Taylor expansion to the total potential energy, we obtain

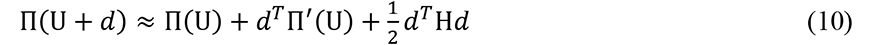

where H is the tangent stiffness matrix and U is the FEM-computed displacement field at the current iteration. The optimal increment *d* that minimizes the right hand side of Eq.(10) is given by

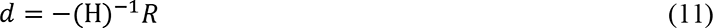

where *R* is the residual force at the current iteration. Then a line search with a positive scalar α is applied to refine the increment and update the displacement field:

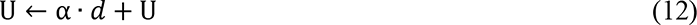

The optimization is considered to be converged if the relative magnitude of the residual force, *R*_*max*_, is close to zero, which is defined by

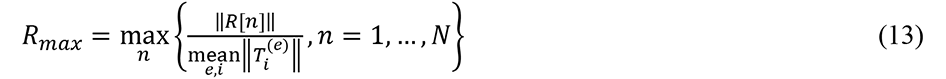

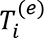 is the nodal equivalent force on the node-i of the element-e. *R*[*n*] is the residual force on the node-n, and *N* is the total number of nodes on an aorta mesh. In our study, we set *R*_*max*_ ≤ 0.005.

In FEA packages, the FE solver usually starts from an equilibrium state (*R* ≈ 0), and the external load increases gradually in a sequence of pseudo time steps. When the external load increases by one small step, the FE solver tries to find the next equilibrium state in several Newton–Raphson iterations using Eq.(11) and Eq.(12). The 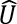 from a DNN is often in a non-equilibrium state. If the FE solver starts from this non-equilibrium state, *d* = −(H)^−1^*R* may point to a completely wrong direction such that NaN or inf will occur in the computing process, resulting in a crash, which is similar to the common issue that a large (time) step in the FE solver leads to a crash. To remedy this issue, we run a L-BFGS [29] based optimization in several iterations to preprocess the 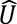 from a DNN, so that the preprocessed 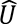 will be closer to an equilibrium state and then can be used as the starting point of the FE solver. L-BFGS approximates the inverse of the matrix H iteratively, and its optimization step can be set to very small (e.g., 1/||*R*||) to prevent NaN and inf in most cases. If the output 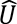 from a DNN has extremely large errors, the L-BFGS based preprocessing stage will crash in the first iteration.

## 2.4 Metrics for performance evaluation by comparing prediction with ground truth

To evaluate the performance of the DNNs, we use the following metrics:

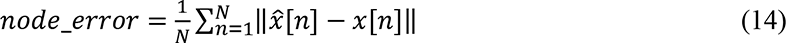

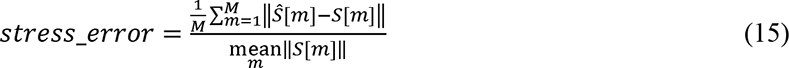

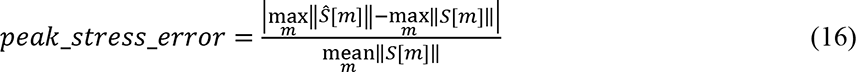

‖∎‖ denotes either scalar absolute value, or vector L2 norm, or matrix Frobenius norm. 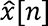 is the predicted position of the node-*n* on the deformed geometry, and *x*[*n*] is the true (FEM-computed) position of the node-*n* on the deformed geometry. *N* is the total number of nodes, and *M* is the total number of elements on an aorta mesh. *S*[*m*] is the true stress at the element-*m* stress, calculated by using Eq.(2) with the true (FEM-computed) deformation. *Ŝ*[*m*] is the stress calculated by using Eq.(2) with the predicted deformation. In the experiments, we use the Von mises stress.

## 2.5 Methods for accuracy assessment of DNNs and DNN-FEM without knowing ground truth

We developed methods to assess the accuracy of the output of a model without knowing the ground truth. It is completely different from performance evaluation in Section 2.4, in which the ground truth is available.

Accuracy assessment of DNNs is very challenging. Out of distribution (OOD) detection (i.e., novelty detection) could assist with such accuracy assessment. If an input sample is OOD (e.g., very different from the training samples), then the output of a model may have large errors. If a sample is considered OOD by a detector, then the model will refuse to make any decision to avoid any disastrous consequences. OOD detection is very challenging, and it has been shown that for image-related applications, many OOD detectors are non-robust [30].

To perform the accuracy assessment of a DNN, a metric is needed to assign a score to the input-output pair of a DNN: a larger score indicates a higher likelihood of having unacceptable output errors. In the experiemnt-1, we tested the following two metrics:

1. Reconstruction error as the score. We built a SSM only using the training samples, and it can be used to reconstruct a sample (i.e., a zero-pressure geometry). The assumption is that if a test sample is very similar to the training samples (i.e., not OOD), then it can be well reconstructed by the SSM, and its displacement field may be well predicted by a DNN model.
2. Average residual force magnitude, 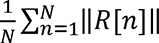, as the score, which is calculated by using the output 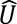 from a DNN. The assumption is that if the output 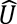 from a DNN is close to an equilibrium state, then it is close to be accurate. We also tried to use *R*_max_ as the score, and it is not better than the average residual force magnitude for DNN accuracy assessment. We note that the value of *R*_max_ determined by the 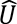 from a DNN is often much higher than 0.005.

We formulate DNN accuracy assessment as a binary classification task: given the output of a DNN for an input sample, a binary classifier will classify the output as accurate or inaccurate based on a classification score (reconstruction error or average residual force magnitude). If the classification score is higher than a threshold (a.k.a. classification threshold), then the output of a DNN is classified as inaccurate, which means the DNN should hand over the task to a FEM-only approach. We use the AUC (area under the ROC Curve) to measure the accuracy of this binary classifier. To create ground-truth class labels, we define that if *peak_stress_error* < *error_threshold*, then the output of a DNN is accurate. *error_threshold* is the maximum acceptable error in peak stress. We choose three values for the *error_threshold*: 0.1, 0.05, and 0.01.

Accuracy assessment of DNN-FEM is much more straightforward, and it can be done by OOD detection. An input of a DNN-FEM integration is considered OOD if any of the two scenarios happens: (1) the L-BFGS based preprocessing step crashes; (2) the number of iterations in the FE solver exceeds a pre-defined number (set to 150 in our experiment). If the output 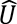 from a DNN has small errors, the FEM-refinement process will complete in a few iterations in a few seconds. If the input is not OOD, the output from DNN-FEM is guaranteed to have acceptable accuracy because we use a strict convergence criterion: *R*_max_ ≤ 0.005. As will be shown in the result section, under this criterion, we obtain *peak_stress_error* ≈ for all test cases.

## 2.6 DNN-FEM integration for solving the inverse problem of heterogeneous material parameter estimation

For biomechanical analysis of huma aorta, there are the following variables: the zero-pressure geometry, the deformed geometry at the systolic phase, and the set of material parameters. The forward problem is to obtain the deformed geometry given the other two as inputs. The inverse problems are (1) obtain the zero-pressure geometry, given the other two as inputs; and (2) obtain the material parameters, given the other two as the inputs. The inverse problem-1 could be solved in a similar way: using a DNN to predict the deformation and using the predicted deformation as the starting point for a FEM-based inverse method. The inverse problem-2 will be straightforward if the material is homogeneous, i.e., every element has the same material parameters. The inverse problem-2 will be very challenging if the material is heterogeneous, e.g., the spatial distribution of the material parameters on the aortic wall is not uniform, which will be the task in this study. For this inverse problem, we developed two methods, a FEM-only inverse method and a DNN-FEM inverse method. The goal of an inverse method is to recover the heterogeneous material parameters for all of the elements, i.e., to obtain {*C*_10_[*m*], *k*_1_[*m*] *k*_2_[*m*], *κ*[*m*] *θ*[*m*], m = 1 to M} a total of 5M variables. In this study, the number of elements M is 4950.

To evaluate the methods, we created a dataset of seven test cases using seven representative zero-pressure geometries and the following material spatial distribution for every case:

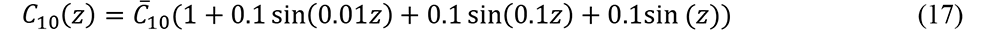

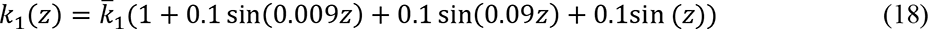

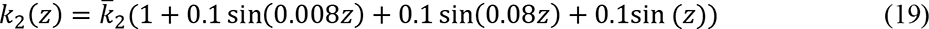

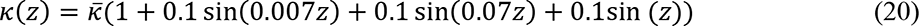

The mean parameters 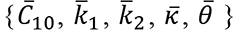 are obtained from our previous studies [31]. The material spatial distribution was created by adding frequency variations to the parameters, except that 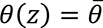 for any z. We used the cylindrical coordinate system, and *z* is the coordinate along the centerline of the aorta. The distribution is visualized by Figure 4 in Section 3.

**Figure 4.**
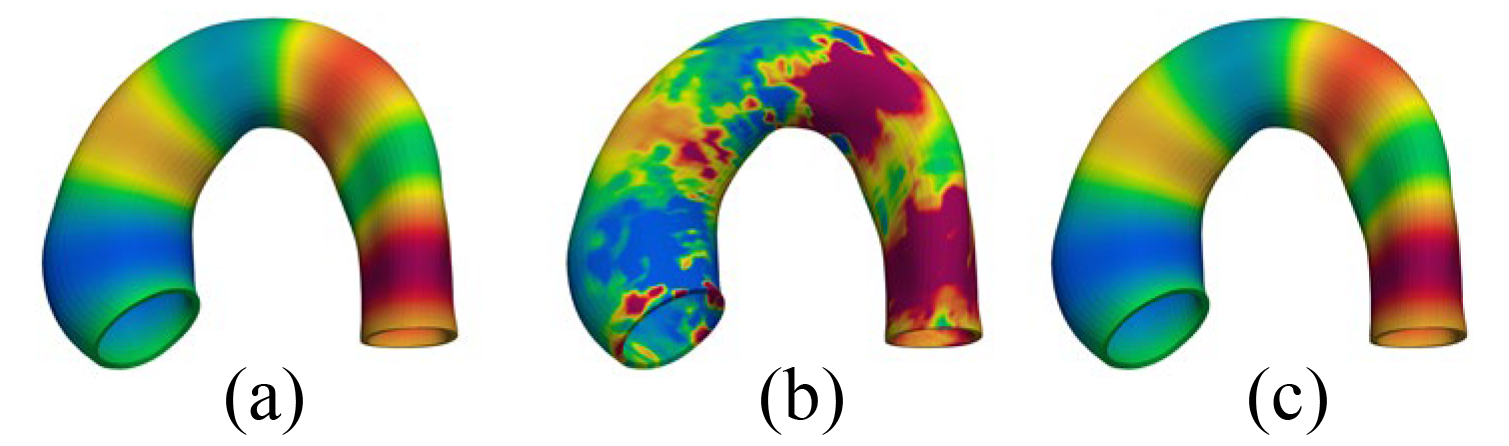
Comparison of the two inverse methods: (a) the ground truth parameter distribution (color-coded), (b) output from the FEM-only inverse method, and (c) output from the DNN-FEM inverse method.

The FEM-only inverse method uses the following loss function:

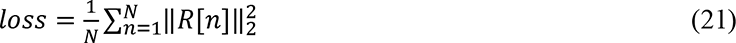

where *R*[*n*] is the residual force on the node-*n*. The loss is a nonlinear function of the 5×4950 unknown material parameters. The best material parameters should lead to the minimum loss at an equilibrium state. Because our FEM is implemented using PyTorch, we use PyTorch automatic differentiation mechanism to automatically obtain the derivative of the loss with respect to each material parameter. Therefore, the optimization can be done in a way similar to training a DNN by backpropagation. We used the L-BFGS optimizer with max iterations of 100000.

The DNN-FEM inverse method uses the same form of the loss function in Eq.(21) with one significant difference: the material parameters of each element are the outputs of an MLP, given by

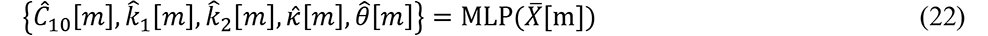

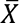[m] is the center of the element-*m* on the mean shape at the zero-pressure state. The MLP is shared by the elements. The MLP poses a constraint on the solution so that the material parameters of the elements are not completely independent. As will be shown in the result section, it helps the optimizer to locate the optimal solution. The internal weights of the MLP become the primary variables to be optimized, and the loss in Eq.(21) becomes a function of the MLP weights. The structure of the MLP will be optimized by a grid search, and the optimal MLP will lead to the minimum loss. We note that the MLP is not pre-trained on any dataset because of the lack of experimental data about spatial distribution of material parameters.

We use the following metric to measure the estimation error of a material parameter:

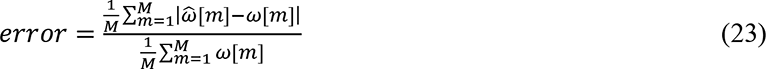

where *ω* represent a true material parameter and 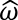 represents the estimation. As will be shown in the result section, the DNN-FEM inverse method performs much better than the FEM-only inverse method.

### 3. Results

## 3.1 Performance evaluation of DNNs by comparing prediction with ground truth

DNN training and testing were conducted on a computer with Nvidia RTX A6000 GPUs. We used the Adamax optimizer with an initial learning rate of 0.001. In the experiment-1, the number of training epochs is 20000 and batch size is 1. In the experiment-2, the number of training epochs is 5000 and batch size is 1.

The performance of the five DNNs for the forward problem is evaluated by comparing their predictions with the ground truth. The metrics in Eq.(14), Eq.(15), and Eq.(16) are used to calculate the prediction errors for each test sample. For the error metrics, we calculate max and average across the test samples. The results of the experiment-1 are shown in Table 1. Since W-Net has the best performance in the exerpiment-1, it is only necessary to use W-Net in the experiment-2, and the result is shown in Table 2. The unit of *node_error* is mm.

**Table 1.** Performance of the five DNNs in the experiment-1 of the forward problem

|  | *node_error*<br>(average) | *node_error*<br>(max) | *stress_error*<br>(average) | *stress_error*<br>(max) | *peak_stress_error*<br>(average) | *peak_stress_error*<br>(max) |
| --- | --- | --- | --- | --- | --- | --- |
| W-Net | **0.0108** | **0.1037** | **1.4624%** | **5.0015%** | **2.9987%** | **51.1089%** |
| TransNet | 0.0192 | 0.2655 | 1.8909% | 9.4478% | 3.7134% | >100% |
| U-Net | 0.0436 | 0.4071 | 2.7889% | 11.8279% | 16.0251% | >100% |
| MeshGraphNet | 0.0696 | 0.5408 | >100% | >100% | >100% | >100% |
| MLP | 0.1295 | 0.7266 | 3.7906% | 11.3773% | 15.5128% | >100% |

**Table 2.** Performance of W-Net in the experiment-2 of the forward problem

| *node_error*<br>(average) | *node_error*<br>(max) | *stress_error*<br>(average) | *stress_error*<br>(max) | *peak_stress_error*<br>(average) | *peak_stress_error*<br>(max) |
| --- | --- | --- | --- | --- | --- |
| 0.0875 | 0.6253 | >100% | >100% | >100% | >100% |

## 3.2 Accuracy assessment of DNNs without knowing ground truth

The two metrics in Section 2.5 are used for DNN accuracy assessment without knowing the ground truth. The results are reported in Table 3, Table 4, and Table 5 with different *error_threshold*. We note that AUC is between 0.5 (equivalent to random guess) and 1 (best). MeshGraphNet has the AUC of 0.5 because its *peak_stress_error* is above the *error_threshold* for almost every test sample.

**Table 3.** AUC of the five DNNs with two different metrics when *error_threshold* = 0.1

|  | W-Net | TransNet | U-Net | MeshGraphNet | MLP |
| --- | --- | --- | --- | --- | --- |
| Reconstruction | 0.3045 | 0.7810 | 0.7417 | 0.5000 | 0.5232 |
| Residual Force | 0.8727 | **0.9293** | 0.8737 | 0.5000 | 0.7971 |

**Table 4.** AUC of the five DNNs with two different metrics when *error_threshold* = 0.05

|  | W-Net | TransNet | U-Net | MeshGraphNet | MLP |
| --- | --- | --- | --- | --- | --- |
| Reconstruction | 0.6202 | 0.7469 | 0.6544 | 0.5000 | 0.5471 |
| Residual Force | 0.8560 | **0.8941** | 0.7226 | 0.5000 | 0.7158 |

**Table 5.** AUC of the five DNNs with two different metrics when *error_threshold* = 0.01

|  | W-Net | TransNet | U-Net | MeshGraphNet | MLP |
| --- | --- | --- | --- | --- | --- |
| Reconstruction | 0.6945 | 0.6945 | 0.6688 | 0.5000 | 0.5425 |
| Residual Force | 0.7573 | 0.7313 | **0.7771** | 0.5000 | 0.7465 |

## 3.3 Evaluation of DNN-FEM for the forward problem

The experiments were performed on a computer with Nvidia Titan V GPUs that support FP64, i.e., double precision. The performance of the five DNN-FEM combinations for the forward problem is evaluated by comparing their outputs with the ground truth. The results are reported in Table 6 and Table 7. Besides the error metrics in Eq.(14), Eq.(15), and Eq.(16), we also compared the time cost of DNN-FEM with the cost of the FEM-only approach. Table 6 and Table 7 show that the average time cost of W-Net-FEM is < 2% of the FEM-only time cost in the two experiments. The “OOD” columns in Table 6 and Table 7 show the number of test samples that are considered as OOD (see Section 2.5).

**Table 6.** Performance of the five DNN-FEM combinations in the experiment-1 of the forward problem

|  | *node_error*<br>(max) | *stress_error*<br>(max) | *peak_stress_error*<br>(max) | time cost<br>(average) | time cost<br>(max) | OOD |
| --- | --- | --- | --- | --- | --- | --- |
| W-Net | 0.0005 | 0.0395% | 0.1056% | 1.6535% | 3.1281% | 0 |
| TransNet | 0.0005 | 0.0393% | 0.0977% | 1.3600% | 3.1090% | 0 |
| U-Net | 0.0005 | 0.0394% | 0.1075% | 1.4737% | 3.5354% | 0 |
| MeshGraphNet | 0.0005 | 0.0393% | 0.1080% | 2.4869% | 7.9622% | 3/137 |
| MLP | 0.0005 | 0.0404% | 0.1081% | 1.9412% | 9.5387% | 0 |

**Table 7.** Performance of the W-Net-FEM in the experiment-2 of the forward problem

| *node_error*<br>(avg) | *node_error*<br>(max) | *stress_error*<br>(max) | *peak_stress_error*<br>(max) | time cost<br>(average) | time cost<br>(max) | OOD |
| --- | --- | --- | --- | --- | --- | --- |
| 0.0002 | 0.0031 | 0.0394% | 0.1116% | 1.4369% | 7.9665% | 61/1781 |

## 3.4 Evaluation of DNN-FEM for the inverse problem

The performance of the two inverse methods is evaluated by comparing their outputs with the ground truth for the seven test cases, using the metric in Eq.(23). The results are reported in Table 8 and Table 9. Figure 4 shows the outputs of the two methods for one of the test cases.

**Table 8.** Performance of the FEM-only inverse method

| case index | $C_{10}$ error | $k_1$ error | $k_2$ error | $\kappa$ error | $\theta$ error |
| --- | --- | --- | --- | --- | --- |
| 1 | 53.2914% | 33.5776% | 48.9675% | 8.7547% | 3.3568% |
| 2 | 37.2363% | 20.0167% | 33.1241% | 4.7770% | 2.0503% |
| 3 | 29.2773% | 16.1704% | 29.3357% | 3.6960% | 1.3987% |
| 4 | 27.0346% | 15.3475% | 24.8842% | 3.6410% | 1.2576% |
| 5 | 28.5631% | 18.6473% | 26.0013% | 4.7736% | 2.2159% |
| 6 | 39.4743% | 25.2475% | 33.2588% | 6.8361% | 3.9024% |
| 7 | 19.3399% | 10.8024% | 18.4389% | 2.1912% | 1.2651% |

**Table 9.** Performance of the DNN-FEM inverse method

| case index | $C_{10}$ error | $k_1$ error | $k_2$ error | $\kappa$ error | $\theta$ error |
| --- | --- | --- | --- | --- | --- |
| 1 | 0.0754% | 0.0482% | 0.2199% | 0.0091% | 0.0048% |
| 2 | 0.4097% | 0.1928% | 0.2290% | 0.0373% | 0.0078% |
| 3 | 0.0656% | 0.0356% | 0.1556% | 0.0070% | 0.0019% |
| 4 | 0.2310% | 0.1235% | 0.2515% | 0.0207% | 0.0070% |
| 5 | 0.1263% | 0.0934% | 0.2303% | 0.0154% | 0.0083% |
| 6 | 0.2083% | 0.1752% | 0.2419% | 0.0311% | 0.0051% |
| 7 | 0.5988% | 0.3744% | 0.3320% | 0.0706% | 0.0119% |

### 4. Discussion

For the forward problem, we developed five types of DNNs to predict deformation. The W-Net (Figure 1) has a brand new structure. The TransNet (Figure 2) uses the Transformer backbone and treats circumferential curves as tokens. The U-Net (Figure 3) treats the circumferential curves of an aorta mesh as a sequence of pseudo points. In the experiment-1 of the forward problem, most of the DNNs perform well with stress error < 5% on average (Table 1). For the application of aortic aneurysm rupture/dissection risk assessment, peak stress is more correlated with the risk [32, 33]. W-Net and TransNet have relatively small peak stress errors on average (<5%, Table 1) but have unacceptably large peak stress errors (>50%, Table 1) for some test samples, which reveals the fundamental issue in ML: due to the pure data-drive nature, an ML model may not generalize well on some new data.

By combining a DNN with FEM for the forward problem, it is guaranteed that the output of DNN-FEM is as accurate as FEM if the input is not an outlier (Table 6 and Table 7) because FEM is used to refine the output of a DNN with a strict convergence criterion. Each DNN-FEM integration runs magnitudes faster than the FEM-only approach because FEM is initialized with DNN output that is very close to the final solution at the systolic phase, and therefore it only takes a small number of iterations to converge to the final solution. In the FEM-only approach, to reach to the final solution, FEM needs to inflate the aorta mesh model from the zero-pressure state to the systolic phase in numerous increments. The combination of W-Net and FEM works the best in the experiment-1, and therefore it is chosen for the experiment-2. In both experiments, the W-Net-FEM integration has peak stress error up to ∼0.1% and only has additional time cost of < 2% on average.

An alternative approach could be performing DNN accuracy assessment without knowing the ground truth, so that a DNN would have the capability of knowing the accuracy of its output. If the output of a DNN is classified to be inaccurate, then the current analysis task may be handed over to the FEM-only approach. If successful, it would be a pure ML approach to guarantee the output accuracy of a DNN if the input is not an outlier. Towards this goal, we developed a binary classification approach to classify the output of a DNN as inaccurate or accurate, by using two different metrics to obtain the classification score: reconstruction error and residual force. We find out that the metric of residual force performs much better than the metric of reconstruction error (Table 3, Table 4, and Table 5). AUC is used to evaluate the binary classification approach. When the maximum acceptable error in peak stress is 10%, TransNet achieves the highest AUC of 0.9293. When the maximum acceptable error in peak stress is reduced to 1%, AUC of TransNet decreases to 0.7313. This experiment shows that DNN accuracy assessment is a promising and challenging direction, not ready to be used for clinical applications.

Uncertainty assessment of DNNs is another approach related to the abovementioned accuracy assessment. Uncertainty includes data uncertainty and model uncertainty. This approach often needs to change learning algorithms or internal structures of DNNs, and therefore it has high complexity [34]. From the perspective of bias and variance estimation, accuracy assessment refers to the estimation of bias (i.e., difference between the prediction and the ground truth), and uncertainty assessment refers to the estimation of variance (i.e., prediction variations), and therefore the two assessments are not the same.

For the inverse problem of identifying heterogeneous material properties, we designed the DNN-FEM inverse method and the FEM-only inverse method, using the same form of loss function. A DNN serves as a regularizer so that the material parameter distribution will not be irregular (Table 8, Table 9, and Figure 4). The Virtual Fields Method (VFM) [35] has been proposed for heterogeneous material parameter identification, and it needs to repeatedly solve the forward problem given an estimation of the material parameters, which is not needed by our method.

This study is different from our previous work using ML models as fast surrogate of FEA [16], which is a data-driven, black-box approach. This study integrates FEM and DNN, for which a DNN is strategically placed in front of FEM to provide an initial solution of the forward problem. Another technical difference is that, in this study, the DNNs predict deformation, and stress is calculated from deformation; in our previous work, the DNN predicts stress only, and it cannot be refined by FEM because the deformation is unknown. Therefore, this study is a significant extension of our previous work.

### 5. Conclusion

We developed new methods to achieve a synergistic integration of DNNs and FEM to overcome each other’s limitations. For the forward problem, DNNs run magnitudes faster than FEM but may have unacceptable errors in the output. The DNN-FEM integration is cable of OOD detection and guarantees the accuracy when the input is not OOD, and it still works magnitudes faster than the FEM-only approach. For the inverse problem, our method uses a DNN as a regularizer to improve the accuracy of material parameter estimation, using the loss function based on the FEM principles.

## Acknowledgement

This work was supported in part by the NIH grant R01HL158829.

## Appendix

## 1. Grid search to optimize DNN structures for the forward problem

We used grid search to optimize the structures of W-Net, TransNet, U-Net, and MeshGraphNet, using the training set and the validation set. After training, the best structure should lead to the best performance on the validation set. Since the MLP encoder-decoder serves as the baseline, we fixed its structure: the encoder is a linear projection layer, and the decoder has five hidden layers with 512 units per layer. Due to limited computation resources, we could only evaluate a small number of structures for the four DNNs. The primary goal of our study is to show DNN and FEM can be combined to improve performance and handle OOD. The seeking of optimal DNN structures is only the secondary goal.

### 1.1 W-Net structure optimization for the experiment-1

Every MLP in W-Net has at least one hidden layer and only one output layer. The hidden dimension of the *MLP*_1_ is fixed to 128, and the number of hidden layers of the *MLP*_1_ is fixed to 3. The number of hidden layers of the *MLP*_2_ is fixed to 2. The number of hidden layers of the *MLP*_3_ can vary. We use sin function as the activation in *MLP*_3_, and use Softplus as the activation for the other MLPs. The hidden dimension of *MLP*_2_ and *MLP*_3_ is the same. The dimension of the code vector is fixed to 3. The grid search result is reported in Table A1. For the experiment-2, we reduce the hidden dimension to 256 in order to reduce training time cost.

**Table A1.**
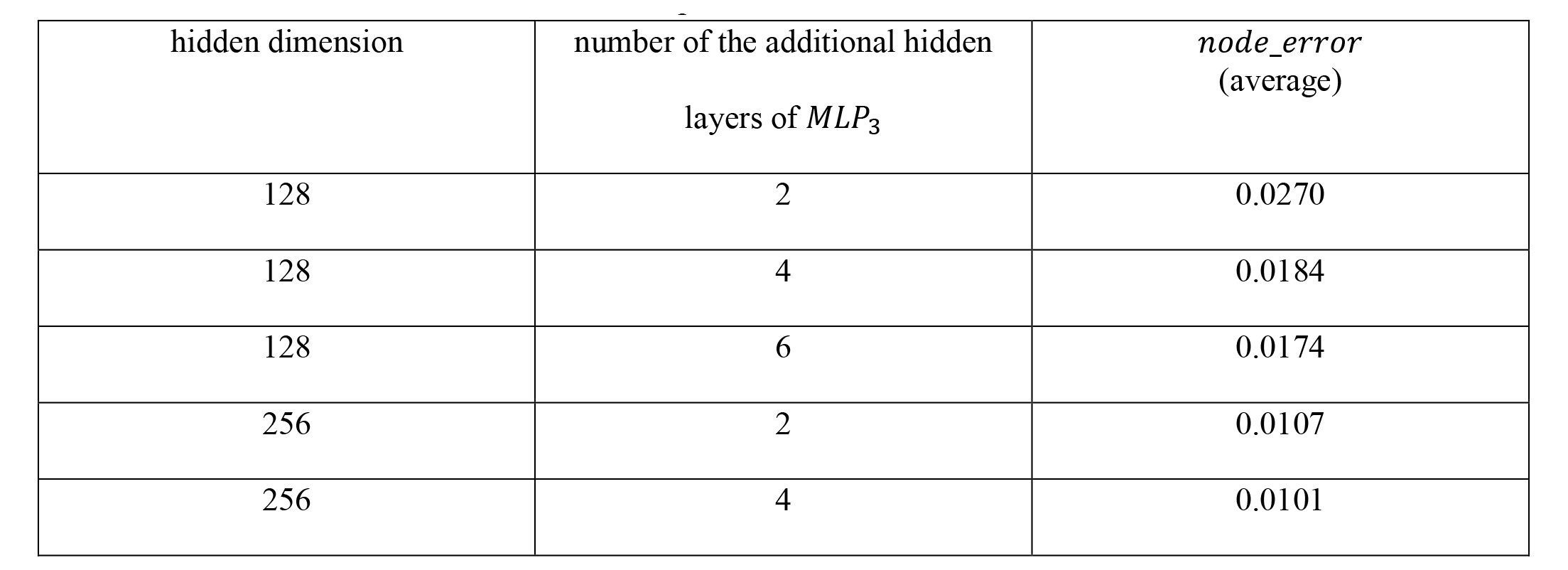
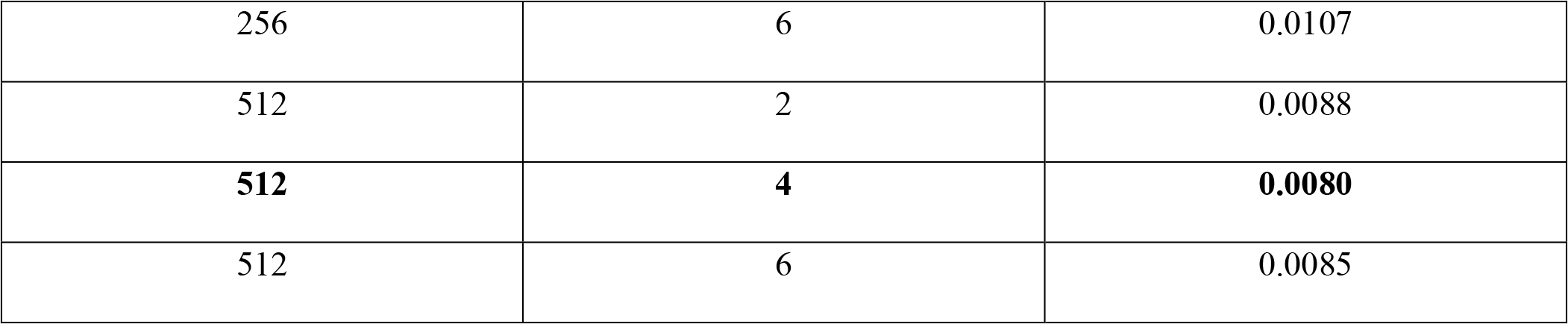
Grid search result of W-Net for the experiment-1

| hidden dimension | number of the additional hidden<br>layers of $MLP_3$ | $node\_error$<br>(average) |
| --- | --- | --- |
| 128 | 2 | 0.0270 |
| 128 | 4 | 0.0184 |
| 128 | 6 | 0.0174 |
| 256 | 2 | 0.0107 |
| 256 | 4 | 0.0101 |
| 256 | 6 | 0.0107 |
| 512 | 2 | 0.0088 |
| **512** | **4** | **0.0080** |
| 512 | 6 | 0.0085 |

### 1.2 TransNet structure optimization

The number of hidden layers in the MLP for position encoding is fixed to 2. The dimension of the hidden layers and the embedding dimensions are kept the same. The grid search result is reported in the Table A3.

**Table A3.** Grid search result of TransNet

| hidden/embedding<br>dimension | number of multi-head<br>self-attention layers | number of heads | *node_error*<br>(average) |
| --- | --- | --- | --- |
| 128 | 4 | 8 | 0.0122 |
| 128 | 4 | 16 | 0.0118 |
| 256 | 1 | 1 | 0.0171 |
| 256 | 1 | 4 | 0.0150 |
| 256 | 1 | 8 | 0.0141 |
| 256 | 1 | 16 | 0.0136 |
| 256 | 2 | 8 | 0.0122 |
| **256** | **2** | **16** | **0.0116** |
| 256 | 4 | 8 | 0.0138 |
| 256 | 4 | 16 | 0.0145 |
| 512 | 4 | 8 | 0.0233 |
| 512 | 4 | 16 | 0.0235 |

### 1.3 U-Net structure optimization

The only structural parameter of the U-Net is the output dimension of the hidden layers, which is denoted by “K” in Figure 3, and it is varied from 64 and 512. The grid search result is reported in Table A4.

**Table A4.** Grid search result of U-Net

| hidden/embedding dimension | *node_error*<br>(average) |
| --- | --- |
| 64 | 0.0639 |
| 128 | 0.0449 |
| 256 | 0.0349 |
| **512** | **0.0317** |

### 1.4 MeshGraphNet structure optimization

Training of the MeshGraphNet is significantly slower than training the other DNNs. Therefore, we only adjusted the number of blocks in the processor and used the recommending settings in the original paper for the other parameters. For example, it took about 7 days to train the model with 15 layers on a Nvidia RTX A6000 GPU. The grid search result is reported in Table A5.

**Table A5.**
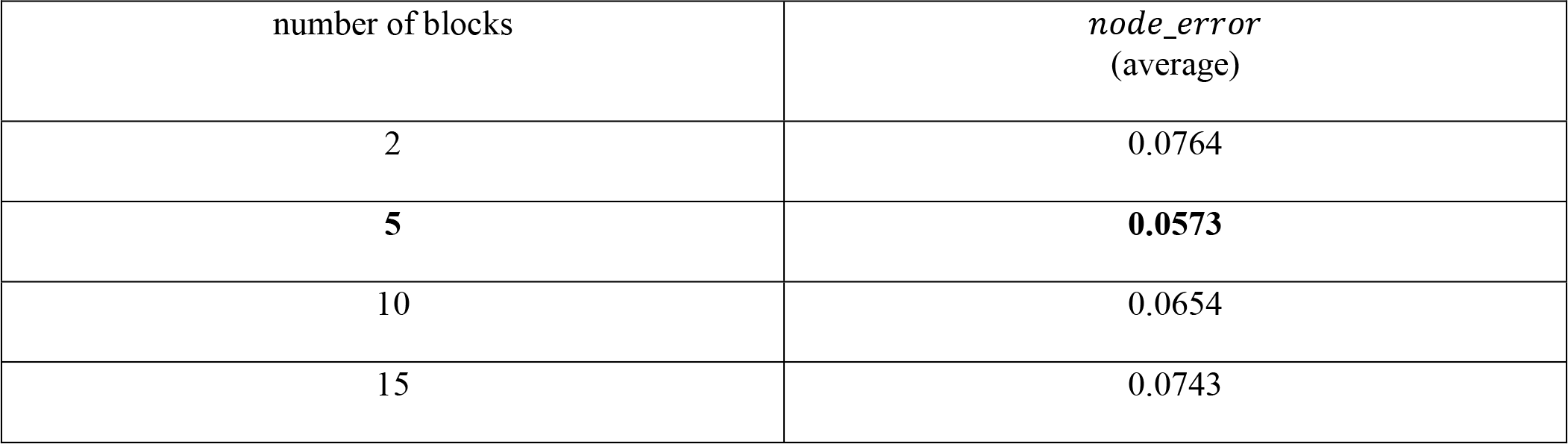
Grid search result of MeshGraphNet

### 2. Grid search to optimize DNN structures for the inverse problem

We used MLP for the inverse problem. We varied the number of layers and the number of units per hidden layer in the grid search. We also compared two activation functions: sin and softplus. The results for one of the test cases are reported in Table A6 and Table A7, which show that sin is much better than softplus.

**Table A6.**
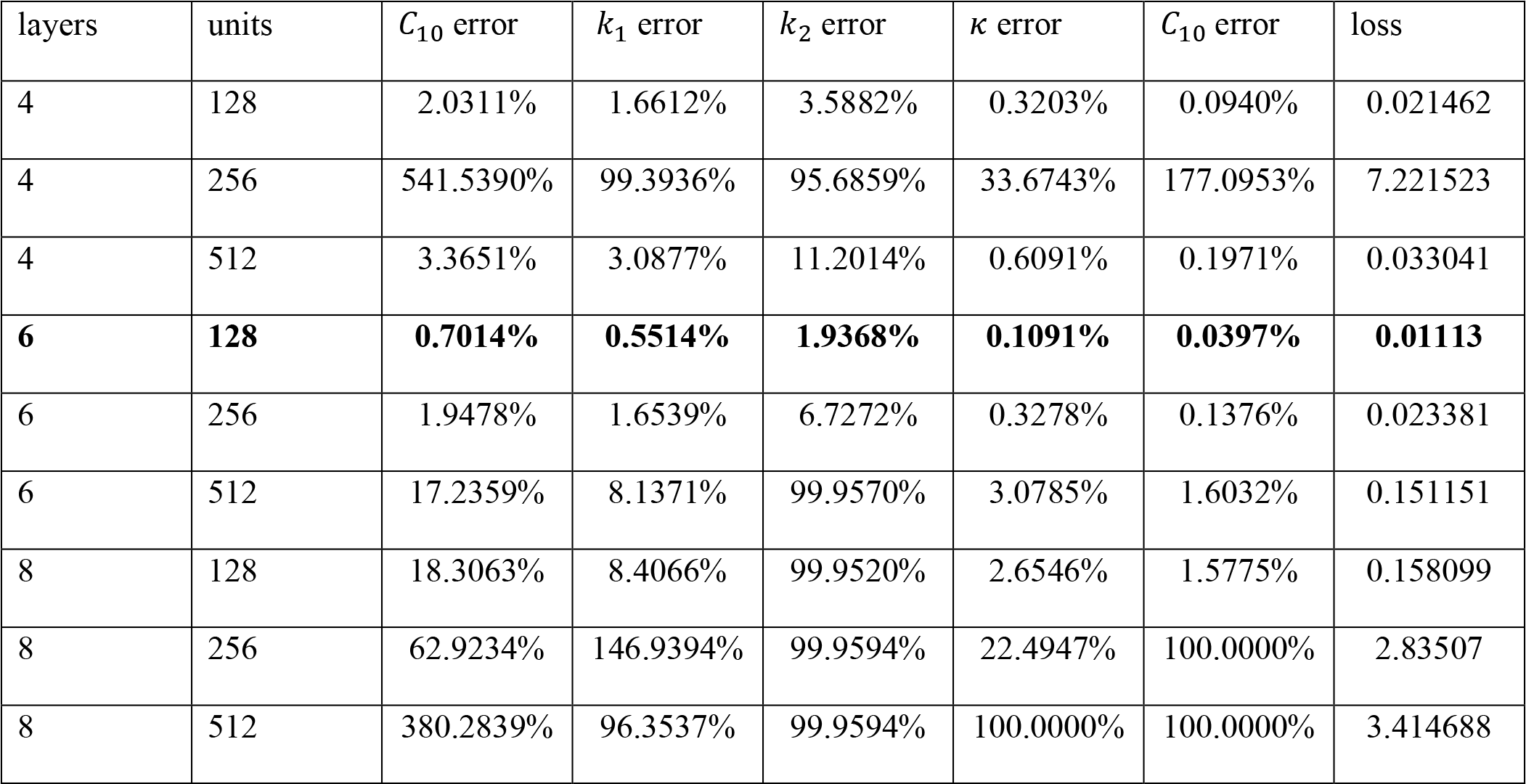
Grid search result using sin activation in MLP for the inverse problem

**Table A7.**
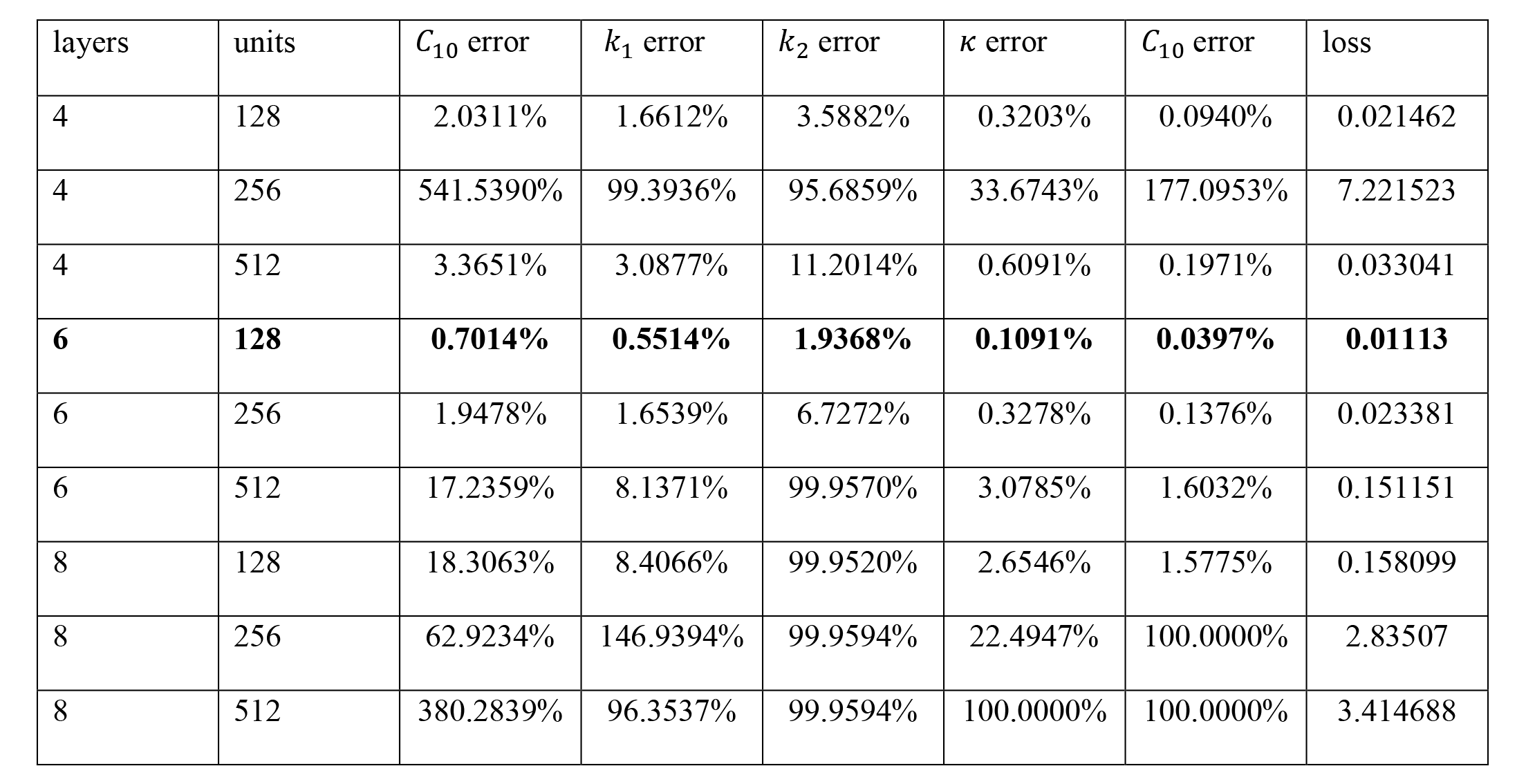
Grid search result using softplus activation in MLP for the inverse problem

